# *FOXP* Genes Regulate Purkinje Cell Diversity in Cerebellar Development and Evolution

**DOI:** 10.1101/2024.11.07.622485

**Authors:** Nagham Khouri-Farah, Qiuxia Guo, Thomas A. Perry, Ryan Dussault, James Y.H. Li

## Abstract

Mammalian cerebellar development is thought to be influenced by distinct Purkinje cell (PC) subtypes. However, the degree of PC heterogeneity and the molecular drivers of this diversity have remained unclear, hindering efforts to manipulate PC diversification and assess its role in cerebellar development. Here, we demonstrate the critical role of *Foxp* genes in cerebellar development by regulating PC diversification. We identified 11 PC subtypes in the embryonic mouse cerebellum through single-cell RNA sequencing. Using a novel unsupervised method, we mapped these subtypes in three-dimensional space, revealing discrete PC subtypes predictive of adult cerebellar organization, including longitudinal stripes and lobules. These subtypes exhibit unique combinations of *Foxp1*, *Foxp2*, and *Foxp4* expression. Deletion of *Foxp2* and *Foxp1* disrupts PC diversification, leading to altered cerebellar patterning, including the loss of a specific *Foxp1*-expressing subtype and the cerebellar hemisphere. The *Foxp1*-expressing PC subtype is much more abundant in the fetal human cerebellum than in mice, but rare in the chick cerebellum, correlating with cerebellar hemisphere size in these species. This highlights the significance of *Foxp1*-expressing PCs in cerebellar hemisphere development and evolution. Therefore, our study identifies *Foxp* genes as key regulators of PC diversity, providing new insights into the developmental and evolutionary underpinnings of the cerebellum.

## INTRODUCTION

Despite its uniform cytoarchitecture, the human cerebellar cortex is partitioned into discrete functional domains as revealed by neuroimaging ^1^. Notably, the cerebellar hemisphere, which is a unique feature in mammals and is greatly enlarged in primates ^2^, participates in cognitive and affective processes ^1^. Alterations in activity originating from the cerebellar hemisphere result in autism spectrum disorder-like behaviors in mice and humans ^3,4^. Currently, our knowledge of the developmental parcellation of the cerebellar cortex and formation of the cerebellar hemisphere is limited.

Anatomical, molecular, and physiological evidence suggests that the mammalian cerebellum is intricately patterned and organized as a series of transverse zones and longitudinal stripes ^5,6^. Transverse fissures divide the cerebellar cortex into lobes and lobules along the anteroposterior (AP) axis, while variations in lobulation delineate three domains from medial to lateral: the medial vermis, paravermis, and hemisphere ^2^. The tripartite division is further subdivided by alternating longitudinal stripes of Purkinje cells (PCs) that express Aldoc or Plcb4 ^5,6^. These parasagittal PC stripes align with afferent terminal fields and efferent projections, serving as a foundation for functional partitioning of the cerebellar cortex ^5,6^. Expression studies using a limited number of markers have revealed molecularly distinct PC subtypes in the developing mouse cerebellum ^7–11^. Heterogeneous PC subtypes can provide a framework for the organization of cerebellar afferent topographies, as they are the earliest-born and sole output neurons of the cerebellar cortex ^5,6^. Although single-cell RNA-seq (scRNA-seq) has the potential to systematically characterize the molecular heterogeneity of PCs, a crucial obstacle is the loss of the spatial context of the technology. This difficulty is exacerbated by complex morphological transformations of the cerebellum during the perinatal stages. Furthermore, it is difficult to relate embryonic PC subtypes to their adult counterparts because of the dynamic expression of markers in PC subpopulations. Finally, we lacked an entry point to specifically perturb PC diversification and to assess its impact on cerebellar development.

In this study, by integrating spatial protein expression analysis and scRNA-seq, we identified 11 molecularly distinct PC subtypes, mapped them in three-dimensional (3D) space, and related them to alternating Aldoc-positive and negative (Aldoc^+/-^) stripes in the adult cerebellum. These subtypes are defined by the combined expression of three *Foxp* genes (*Foxp1*, *Foxp2*, and *Foxp4*). Loss-of-function studies have demonstrated that *Foxp1* and *Foxp2* are essential for PC diversification. Their removal resulted in progressive reduction and eventual loss of the cerebellar hemispheres, coinciding with disruption of a specific PC subtype. Remarkably, this subtype is significantly expanded in the human cerebellum and largely absent in the chicken cerebellum. Taken together, our findings show that PC diversification controls pattern formation in the cerebellar cortex, and contributes to cerebellar evolution.

## RESULTS

### Identification of 11 molecularly distinct Purkinje cell subtypes in the embryonic mouse cerebellum

PCs are produced between embryonic days (E) 10 and E13 in mice ^12^. We have previously described at least five molecularly distinct PC subtypes at E13.5, indicating that PC heterogeneity is established soon after PCs are born ^13^. To characterize PC heterogeneity at the later embryonic stage, we performed scRNA-seq of the mouse cerebella at E16.5 and E18.5. We assigned identity to each cell cluster from the combined dataset according to known cell type-specific markers, recovering all major cerebellar cell types (Fig. 1A, Supplementary Table S1). The molecular features of individual cell types were remarkably consistent between E16.5 and E18.5 (Fig. S1A), in agreement with our previous finding that the differentiation of cerebellar cell types becomes subtle and gradual toward the end of gestation ^5^.

**Figure 1.**
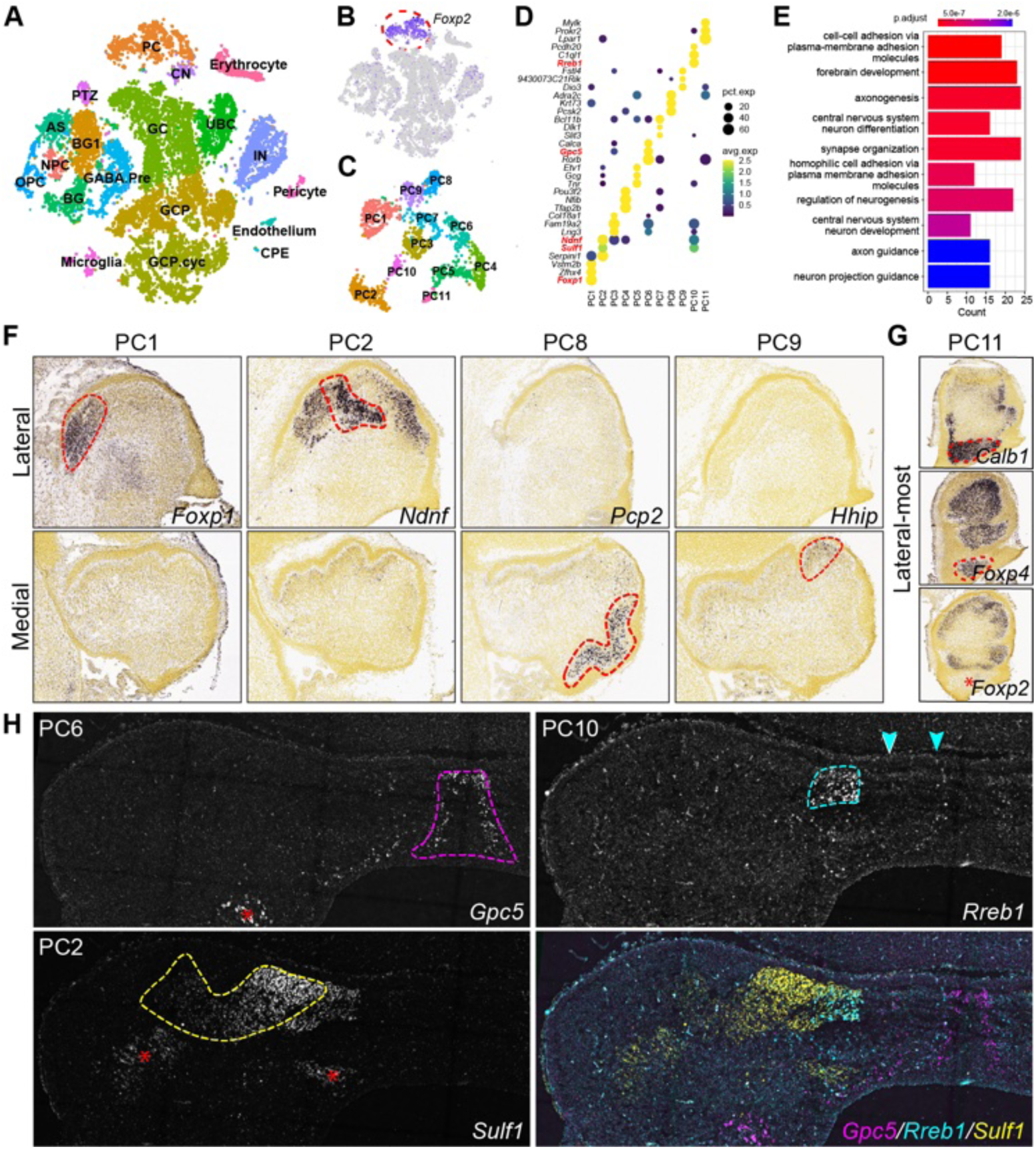
scRNA-seq identification of 11 PC subtypes in the mouse cerebellum before birth. (**A**) Visualization of major cerebellar cell types using t-distributed stochastic neighbor embedding. Dots denote cells, and colors represent cell types. Abbreviations: AS, astrocytes; BG/BG1, Bergmann glia; CN, cerebellar nuclear neurons; CPE, choroid plexus epithelium; GABA.Pre, GABAergic neuron precursor; GC, cerebellar granule cell; GCP, granule cell progenitor; GCP.cyc, proliferative granule cell progenitor; IN, interneuron; NPC, neural progenitor cell; OPC, oligodendrocyte progenitor cell; PC, Purkinje cell; PTZ, posterior transitory zone; UBC, unipolar brush cell. (**B**) Selection of PCs based on *Foxp2* expression. (**C**) Visualization of the 11 PC subtypes using uniform manifold approximation and projection (UMAP). (**D**) Dotplot showing the molecular features of the PC subtypes. The markers in red were validated using expression data. (**E**) Functional enrichment of genes that distinguish different PC subtypes. (**F** and **G**) Region-specific expression of markers for five PC subtypes in sagittal sections of E18.5 mouse cerebella (from the Allen Developing Mouse Brain Atlas). The dashed lines indicate the expression domains in either the medial or the lateral sections. Lateral-most sections in G show that PC11 cells express *Calb1* and *Foxp4*, but lack *Foxp2* transcripts (indicated by an asterisk) in the prospective flocculus. (**H**) RNAscope fluorescence in situ hybridization of coronal sections of the cerebellum at E16.5. The dashed lines outline PCs that are positive for the specific probe. Arrowheads indicate longitudinal stripes of *Rreb1*+ PCs.

We reiterated the cell clustering of PCs recognized by the pan-PC marker *Foxp2* ^13,14^ and identified 11 PC subtypes, provisionally named PC1-11, based on their relative abundance (Fig. 1B-D, Fig. S1B, and Supplementary Table S2). The salient molecular features distinguishing the PC subtypes were cell surface adhesion molecules, including eight members of the cadherin superfamily (Fig. 1E, Fig. S1C and Supplementary Table S2). To evaluate the robustness of the analysis, we re-analyzed two published scRNA-seq datasets of the E16-E18 mouse cerebella ^15,16^ and identified similar heterogeneity and abundance of PC subtypes (Fig. S1D-F).

Based on the Allen Developing Mouse Brain Atlas and fluorescence in situ hybridization (FISH), we approximated the endogenous positions of seven PC subtypes in the embryonic cerebellum (Fig. 1F-H), revealing that these subtypes form discrete clusters. For instance, PC1 cells, characterized by *Foxp1* expression, were confined to the lateral-most region of the cerebellum, whereas PC11 cells, largely lacking *Foxp2* transcripts, were restricted to the flocculonodular lobe (Fig. 1F, G, and Fig. S2). Our findings indicate the presence of at least 11 molecularly distinct PC subtypes with regional specificity in the mouse cerebellum before birth.

### Mapping PC subtypes to their native space using an unsupervised strategy

Assigning all 11 PC subtypes to their original spatial contexts using conventional expression analysis was challenging because each subtype was mostly defined by combinatorial markers. We attempted to develop an objective, autonomous method to map cell clusters identified by scRNA-seq to the 3D space in the cerebellum. Through single-cell analysis of the transcriptome and chromatin landscape, we previously identified the gene regulatory network and its driver genes in various cerebellar cell lineages ^13,17^. We hypothesized that, once spatially resolved, these driver genes could act as anchors to pinpoint their regulated gene networks within their native spatial context. This approach would enable the assignment of cell clusters identified by scRNA-seq to their original locations in the cerebellum. To accomplish this, we developed a method using cyclic immunofluorescence (CycIF) to profile the expression of 30–50 proteins in serial cerebellar sections at single-cell resolution (Fig. 2A). By combining single-cell protein spatial profiling with scRNA-seq data, we inferred the identity of individual cells within the sections (Fig. 2A). We then aligned consecutive sections based on the shared cellular features ^18,19^ and reconstructed the 3D structure (Fig. 2B). We termed this strategy <u>s</u>ingle-<u>c</u>ell <u>A</u>nchoring <u>N</u>etwork of <u>K</u>ey <u>R</u>egulators to <u>S</u>pace (scANKRS).

**Figure 2.**
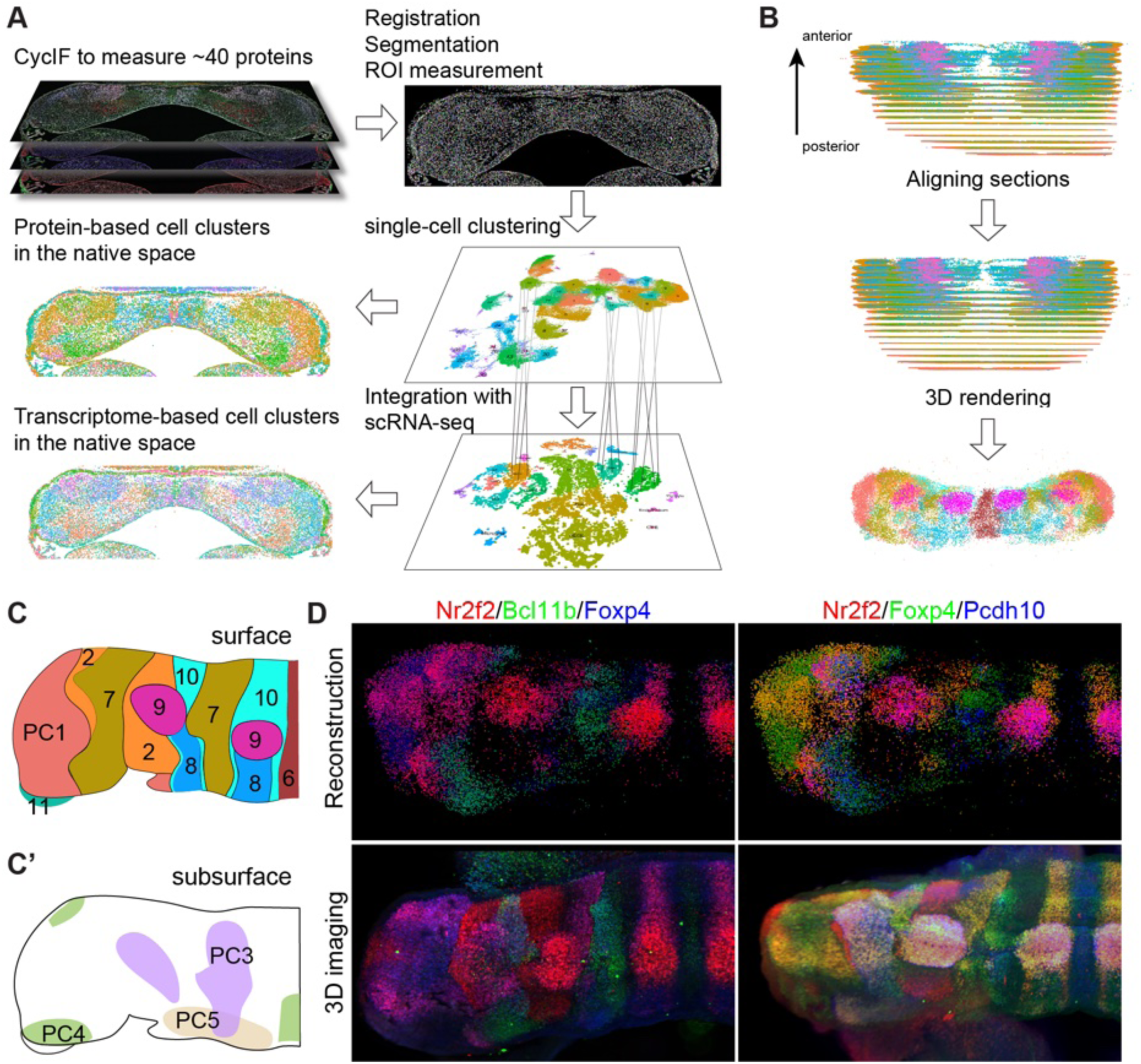
Unsupervised mapping of PC subtypes to a 3D space. (**A**) Schematic representation of the scANKRS workflow that integrates spatial protein profiling and scRNA-seq to assign 11 PC subtypes to serial coronal sections. (**B**) Alignment of serial sections to reconstruct the 3D cerebellar structures. (**C** and **C’**) Illustration showing the spatial distribution of the 11 PC subtypes in the surface (C) and subsurface (C’) in the E17.5 mouse cerebellum, with subtype identities indicated by numbers. (**D**) Comparison of 3D reconstruction with light-sheet fluorescent images of wholemount cerebella following immunofluorescence and tissue clearing.

Using scANKRS, we determined the topographic distribution of the 11 PC subtypes identified by scRNA-seq in the embryonic mouse cerebellum (Fig. 2B). While most PC-subtype clusters appeared near the cerebellar surface, PC3, 4, and 5 were positioned beneath the surface, close to the ventricular zone at E17.5, potentially representing transient-state PCs (Fig. 2C and C’). We confirmed the reconstructed spatial configuration of the PC subtypes through volume imaging using light-sheet fluorescence microscopy (LSFM), following immunolabeling and tissue clearing (Fig. 2D).

To further validate scANKRS, we transferred the transcriptome to individual cells within sections. As expected, the imputed expression patterns of the mRNAs closely matched those of their corresponding protein products (Fig. S3A). We confirmed the accuracy of the imputed spatial expression using FISH ^20^ for the selected molecules (Fig. S3B). The results from different independent experiments were consistent and reproducible (Fig. S3C and D). Thus, by integrating scANKRS with 3D reconstruction and volume imaging, we reliably determined the 3D positioning of the PC subtypes within the embryonic mouse cerebellum.

### The heterogeneity of embryonic PCs predicts their regional specificity in the adult cerebellum

The mammalian cerebellum exhibits an evolutionarily conserved pattern of alternating longitudinal stripes of Aldoc^+^ or Aldoc^−^ PCs ^5^. Single-nucleus transcriptome profiling has confirmed these two main PC groups and their associated molecular features in the adult mouse cerebellum ^21,22^. It also revealed significant regional specificities among the PCs, with the greatest diversity found in the posterior lobules ^22^. By aligning the embryonic and adult scRNA-seq datasets, we found that most of the 11 PC subtypes corresponded to either Aldoc^+^ or Aldoc^−^ PCs (Fig. 3A, B). When the inferred Aldoc identity was projected in the 3D reconstructed cerebellum according to the corresponding PC subtypes, the Aldoc^+^ and Aldoc^−^ precursors formed alternating longitudinal stripes, mirroring the adult pattern (Fig. 3C). Furthermore, aligning adult regional PC identities with embryonic PCs showed that different PC subtypes were associated with specific lobules (Fig. 3D). For instance, PC1 was enriched in the ansiform Crus I and II lobules (CrI and CrII), paramedial lobules (Prm), and paraflocculus, while PC11 was restricted to the flocculus (Fig. 3D).

**Figure 3.**
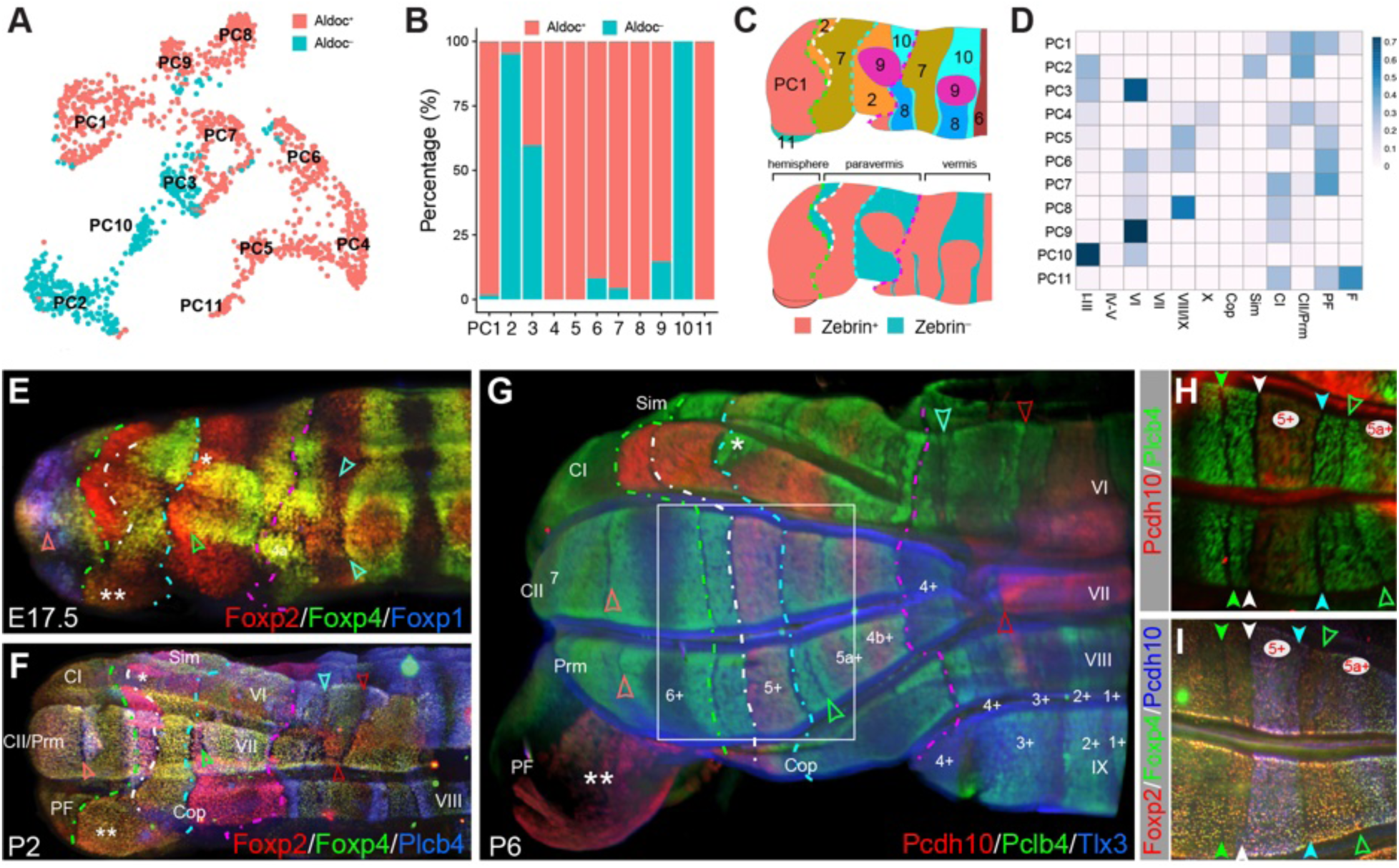
Alignment of embryonic PC subtypes with adult PC phenotypes. (**A** and **B**) UMAP and barplot showing the alignment of embryonic PC-subtype clusters with Aldoc^+^ and Aldoc^−^ PC identities in the adult cerebellum. (**C**) Schematic presentation of PC-subtype clusters (top) and predicted Aldoc^+/-^ phenotypes (bottom) in E17.5 mouse cerebellum. Dashed lines indicate four major PC-sparse gaps. (**D**) Heatmap of the confusion matrix showing the relationship between the PC subtypes and region-specific PCs in the adult cerebellum. (**E-I**) LSFM images depicting the arrangement of PC subtypes in the mouse cerebellum from E17.5 to P6. Colored dashed lines highlight persistent PC-sparse gaps along the AP axis; arrowheads indicate emerging, additional cell-sparse gaps that emerge in the perinatal mouse cerebellum. The single asterisk denotes a triangular landmark that persists from E17.5 and P6, while the double asterisk denotes Pcdh10^+^ PC7 cells in the posterior part of the paraflocculus. Note that Pcdh10 is absent in the second PC7 band in the vermis, adjacent to the magenta dashed line. The boxed area in G is enlarged in H and I; arrowheads indicate the PC-sparse gaps with color matching the dashed lines in G. Aldoc^+^ PC stripes are named as numerals and letters followed by “+”. Abbreviations: Cop, copula pyramidis; CI/CII, crus I and II lobules; F, flocculus; Prm, paramedian lobule; PF, paraflocculus; Sim, simplex lobule

To confirm the relationship between PC-subtype clusters and longitudinal stripes, we performed 3D imaging of immunolabeled cerebella from E14.5 to P6. We found that many PC-subtype clusters and stripes were separated by PC-sparse gaps, visible by immunostaining of nuclear (Foxp proteins) or cytoplasmic (Pchd10 and Plcb4) proteins, indicating that these gaps separate both the soma and processes of PCs (Fig. 3E-I and Supplementary Movie 1). Four major PC-sparse gaps extended along the entire AP axis, persisting from E14.5 to P6, while several minor gaps emerged between E17.5 and P6 (Fig. 3E-I and Supplemental Movie 1). Among the major gaps, the lateral-most gap separated PC1 from the other subtypes, while the medial-most gap was constituted by the boundaries of several PC subtypes (Fig. 3E-G). We propose that these two PC-sparse gaps delineate the junctions between the hemisphere and paravermis, and between the paravermis and vermis (see below). The other two major gaps were along the border between PC7 and PC2 in the paravermis, revealing that PC7 contributes to the 5+ Aldoc stripe in lobule VII, the Pcdh10-positive domain in Crus I, and the posterior paraflocculus (Fig. 3E-I). Therefore, PC-sparse gaps, potentially representing lineage-restriction boundaries, serve as reliable landmarks for linking PC subtypes from the embryonic to postnatal cerebellum. In summary, by correlating molecular features with cellular landmarks, we relate embryonic PC subtype-specific clusters to longitudinal PC stripes in the adult cerebellum.

### Foxp1 and Foxp2 control the development and function of the mouse cerebellum

Different PC subtypes showed distinct combinatorial expression patterns of Foxp1, Foxp2, and Foxp4, which belong to a subfamily of forkhead-box transcription factors (Fig. 4A and B). Previous studies have demonstrated that a global knockout of *Foxp2* causes reduced cerebellar size and foliation in mice ^23–25^, although the underlying mechanism remains unclear. The role of *Foxp1* in cerebellar development remains unclear. We hypothesized that *Foxp1* and *Foxp2* are involved in the regulation of PC diversification, which, in turn, controls cerebellar development. To test this hypothesis, we performed a conditional knockout (cKO) using a floxed (F) allele of *Foxp1* and *Foxp2* in combination with an *En1^Cre^* driver, deleting *Foxp1* and *Foxp2* in the neural plate of the cerebellum and midbrain as early as E8.5. ^26^.

**Figure 4.**
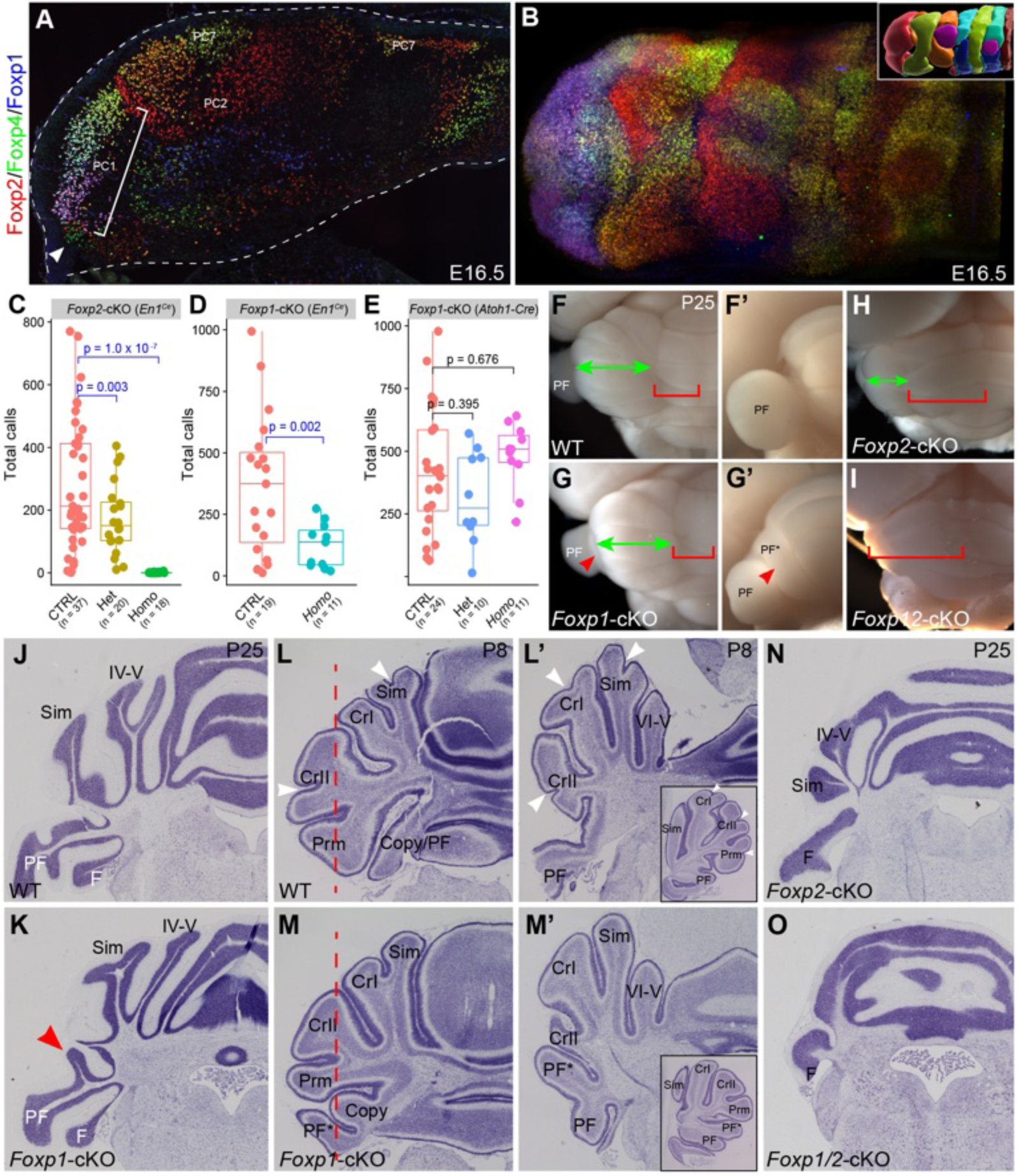
*Foxp1* and *Foxp2* are essential for ultrasonic vocalization and cerebellar development in mice. **(A** and **B)** Immunofluorescence for Foxp1, Foxp2, and Foxp4 in coronal sections (A) and whole mounts (B) of the E16.5 cerebellum. (**C-E**) Boxplots of ultrasonic vocalization (USV) calls from neonatal mice at P6. Horizontal lines within the box denote median values, the lower and upper hinges of the boxplots correspond to the 25^th^ and 75^th^ percentiles, respectively, and dots represent individual samples (n = 37, 20, and 18 in C; n = 19 and 11 in D; n = 24, 10, and 11 in E). P-values in C (*F*_(2, 72)_ = 19.79, *p* = 1.42e-07) and E (*F*_(2, 42)_ = 1.71, *p* = 0.192) were calculated using ANOVA and Tukey’s HSD post-hoc test; the p-value in D (*t*(23.56) = 3.47, *p* = 0.002, 95% CI [98.66, 388.84]) was calculated using two-sided unpaired t-test with Welch’s correction. (**F-I**) Dorsal and side views (F’ and G’) of the P25 cerebellum. Green double arrows indicate the cerebellar hemisphere and paravermis, red brackets demarcate the vermis, and arrowheads denote supernumerary folia of the paraflocculus (PF). (**J-O**) Nissl histology of the P25 (J, K, N, and O) and P8 (L-M’) cerebella. The red arrowhead indicates the extra folia, the white arrowhead denotes the rudimentary fissure present in the WT but not in *the Foxp1*-cKO cerebellum, the asterisk shows the enlarged flocculonodular lobe, and insets in (L’ and M’) show a sagittal section at the level indicated by the dashed line in (L and M).

Unlike global knockouts of *Foxp1* and *Foxp2*, which cause mortality around E14.5 and P15, mice with *Foxp1* and *Foxp2* deletions in the *En1* lineage survived. It has been shown that global *Foxp2* knockout eliminates ultrasonic vocalizations (USVs) in pups separated from their mothers ^24,25,27^. *En1^cre^*-mediated deletion of either a single allele or both alleles of *Foxp2* or deletion of *Foxp1* significantly reduced USVs in mice (Fig. 4C and D) ^28^. *Foxp1* is also expressed in glutamatergic neurons within the cerebellar nuclei ^13^. However, specific removal of *Foxp1* from glutamatergic neurons using an *Atoh1-Cre* transgene ^29^, did not affect USVs, suggesting that *Foxp1* regulates USVs through its action in PCs rather than in the cerebellar nuclei (Fig. 4E). Our findings indicate that *Foxp1* and *Foxp2* are crucial for cerebellar function in the regulation of USVs in mice.

We next examined cerebellar morphology and histology in *Foxp1/2* mutant mice between P21 and P23, when cerebellar morphogenesis is mostly complete ^30^. Although overall cerebellar foliation was preserved in *Foxp1*-cKO mice, the paraflocculus exhibited an abnormal branch, displacing parts of the CrI, CrII, and Prm lobules (n ≥ 3; Fig. 4F-G’, J, and K). The *Foxp1*-cKO phenotype was apparent at P6 (Fig. 4L-M’ and Fig. S4A), highlighting the critical role of *Foxp1* in the formation of the cerebellar hemisphere.

Loss of *Foxp2* resulted in a relatively normal cerebellar vermis, although the size of the central lobe was markedly reduced (Fig. S4B). The cerebellar hemisphere, together with the paraflocullus, was diminished in *Foxp2*-cKO mice and nearly absent in *Foxp1/2* double conditional knockout (*Foxp1/2*-cKO) mice (Fig. 4H, I, N, O, Fig. S4B, and S4C). In *Foxp2*-deficient mice, the flocculonodular lobe was notably enlarged, forming a continuous folium rather than a distinct lobe X in the vermis and flocculus laterally, as observed in wild-type (WT) mice (Fig. S4C). Volumetric measurements showed that the flocculus was significantly larger in *Foxp2*-cKO mice than that in littermate controls at E17.5 (Fig. S4D). We confirmed differential changes in the paraflocculus and flocculus due to the loss of *Foxp1* and *Foxp2* by immunofluorescence (IF) for Hspb1 to mark PCs and Tlx3 to mark granule cells (GCs) in the paraflocculus (Fig. S4E). Despite these extensive morphological defects, cortical lamination and cytoarchitecture were remarkably intact in the cerebella lacking *Foxp1* and *Foxp2* (Fig. 4J-O and Fig. S4B-C). Thus, *Foxp1* and *Foxp2* are essential regulators of regional and lobular identity in the developing mouse cerebellum.

### Specific deletion of the cerebellar hemisphere in the absence of Foxp1 and Foxp2

The markedly reduced lateral part of the *Foxp2*-deficient cerebellum suggests loss of the cerebellar hemisphere. However, since the hemisphere is defined by a unique set of gyri and fissures, we sought to determine whether *Foxp2* deletion disrupted its identity rather than altering its morphology. PCs in the cerebellar hemisphere are primarily innervated by climbing fibers from the principal olive (PO) and dorsal accessory olive (DAO), and two subnuclei of the inferior olive in the medulla oblongata ^5,6^. Given that neuronal survival generally relies on the correct targeting of connections ^31^, we hypothesized that the loss of hemispheric identity would lead to a loss of PO and DAO neurons. Histology and *Foxp2* expression in the inferior olive were indistinguishable between *Foxp2*-cKO mice and their WT littermates at P2, indicating that the initial formation of the inferior olive and *Foxp2* expression was unaffected (Fig. 5A and B).

**Figure 5.**
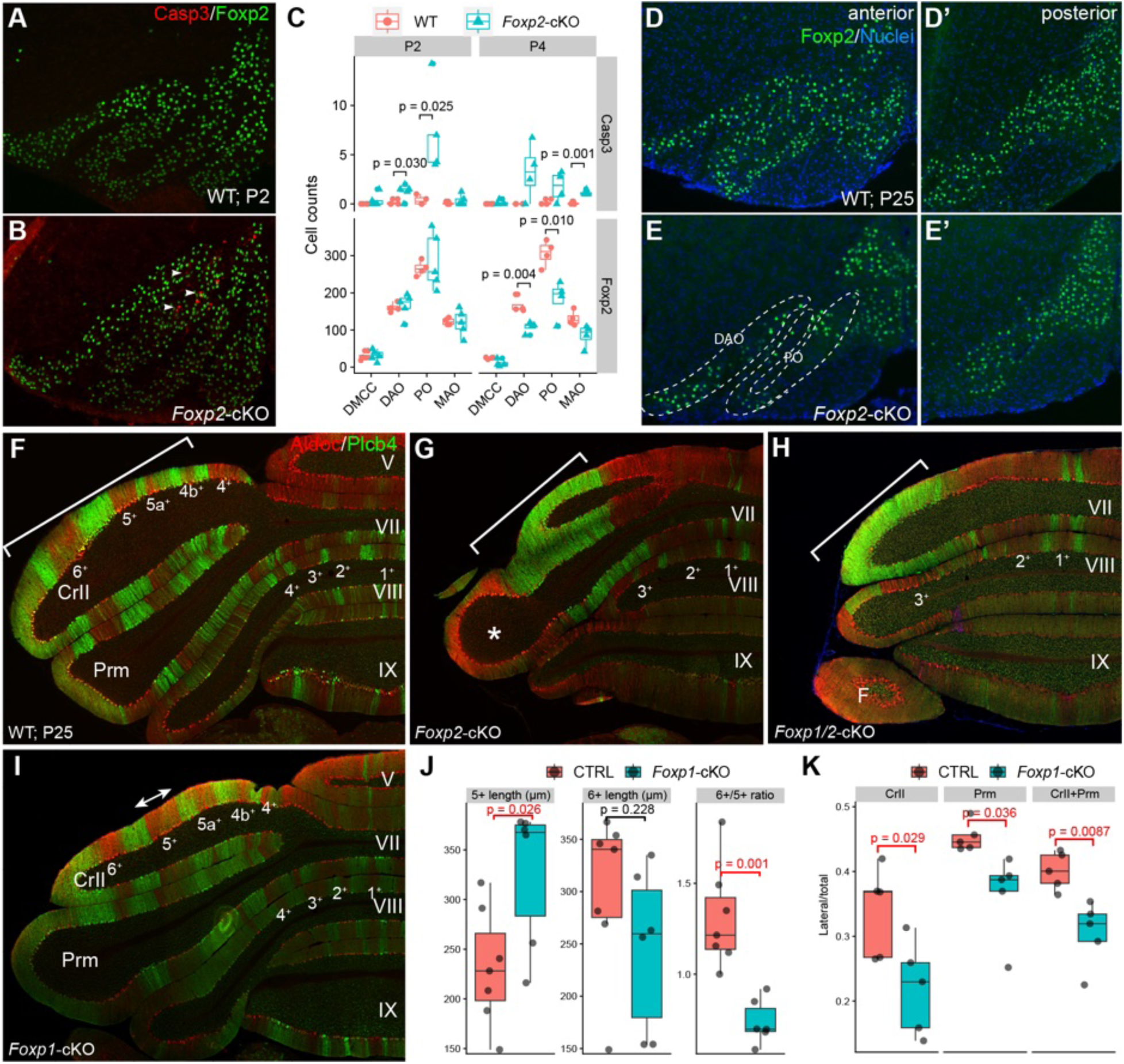
Interactions between *Foxp1* and *Foxp2* control cerebellar hemisphere development. (**A** and **B**) Immunofluorescence showing apoptotic cells (activated Casp3) and inferior olive neurons (Foxp2). Arrowheads denote apoptotic cells in principal olives (PO). (**C**) Quantification of Casp3^+^ apoptotic cells and Foxp2^+^ neurons in different inferior olivary nuclei. (**D-E’**) Immunofluorescence indicates that in *Foxp2*-cKO mice at P25, the PO and dorsal accessory olive (DAO) are absent, while other subnuclei remain largely intact, especially those in the posterior part of the inferior olive. (**F-I**) Immunofluorescence for Aldoc and Plcb4 in coronal sections of the P25 cerebella. Brackets demarcate the alternating Aldoc+ and Aldoc-stripes lateral to the vermis in wildtype (WT) and *Foxp1*-cKO (P1-cKO) cerebella, and the single Aldoc^-^/Plcb4^+^ band in *Foxp2*-deficient cerebella. The asterisk (G) indicates the residual cerebellar hemisphere. (**J, K**) Quantitative analyses of the width of Aldoc+ stripe 5+ and 6+ and the relative area of ansiform crus II (CrII) and paramedial (Prm) lobules lateral to stripe 5-. The respective p-values were calculated by two-sided unpaired t-test with Welch’s correction in (C, J, and K). Abbreviations: DMCC, dorsal medial cell column; MAO, medial accessory olive; F, flocculus; PF, paraflocculus; Sim, simplex lobule.

However, a significant increase in apoptotic cells, marked by activated caspase 3 (Casp3), was observed in the PO and DAO at P2, with a marked reduction in PO and DAO neurons labeled by Foxp2 immunoreactivity at P4 in *Foxp2*-cKO mice (Fig. 5C). By P25, PO and DAO were diminished in *Foxp2*-cKO mice, whereas other inferior olivary subnuclei were relatively intact (Fig. 5D-E’). Therefore, we show that the inferior olivary subnuclei targeting the cerebellar hemisphere are selectively affected in *Foxp2*-cKO mice, supporting the notion that *Foxp2* is essential for cerebellar hemisphere formation.

To further corroborate the loss of the hemisphere in *Foxp2-*deficient cerebella, we analyzed ML patterning of the cerebellum using IF for Aldoc and Plcb4. The vermis shows a characteristic arrangement of four Aldoc^+^/Plcb4^−^ stripes (1+, 2+, 3+, and 4+) interspersed with Aldoc^−^/Plcb4^+^ stripes (1-, 2-, and 3-) in lobules VII and VIII on either side of the midline (Fig. 5F). Flanking the vermis, the paravermis and hemisphere display six alternating Aldoc^+^/Plcb4^−^ stripes (4+, 4b+, 5a+, 5+, 6+, and 7+) and Aldoc^−^/Plcb4^+^ stripes in CrII and Prm lobes (Fig. 5F). In *Foxp2*-deficient mice, although the alternating Aldoc^+^/Plcb4^−^ and Aldoc^−^/Plcb4^+^ stripes persisted in lobules VII and VIII, the 3-stripe was either reduced or shifted laterally, creating a broader Aldoc^+^/Plcb4^−^ domain in the cerebellar vermis (Fig. 5G and H). Notably, the alternating Aldoc^+^ and Aldoc^−^ stripes lateral to the vermis were replaced by a broad Aldoc^−^/Plcb4^+^ domain in both *Foxp2*-cKO and *Foxp1/2*-cKO cerebella (Fig. 5G and H). An Aldoc^+^ region, likely representing the residual hemisphere, was observed in the lateral-most area of *Foxp2*-cKO cerebella but was absent in *Foxp1/2*-cKO cerebella, supporting the stepwise reduction of the cerebellar hemisphere in these mutants (Fig. 5G and H).

*Foxp1*-cKO mice exhibited subtitle defects mainly in the lateral cerebellum. Stripe 5+ was significantly widened and became broader than stripe 6+, reversing the pattern observed in littermate controls (Fig. 5I and J). When normalized to the total area, the region lateral to stripe 5-in lobules CrII and Prm was significantly reduced in *the Foxp1*-cKO cerebella at P25 (Fig. 5I and K). These findings indicate that *Foxp1* deletion selectively impairs the development of the lateral-most region of the cerebellum, consistent with its restricted expression in the cerebellar anlage.

In summary, our data demonstrate that *Foxp2*, together with *Foxp1*, is essential for the development of the cerebellar hemisphere.

### Foxp1 *and* Foxp2 concertedly regulate PC diversification

To test our hypothesis that *Foxp1* and *Foxp2* regulate cerebellar development by controlling PC diversification, we quantified the abundance of various cerebellar cell types in *Foxp1*-cKO, *Foxp2*-cKO, *Foxp1/2*-cKO, and their littermate controls at E16.5 with scANKRS. We found no significant differences in the relative abundance of the major cell types among the four genotypes (Fig. S5A). The total number and proportion of PCs in individual coronal sections at the corresponding AP positions were similar between the WT and *Foxp2*-deficient cerebella, with only a slight decrease at the two most anterior positions (Fig. S5B and C; p > 0.05; two-sided unpaired t-test with Welch’s correction). In contrast, the composition of different PC subtypes was markedly altered by the loss of *Foxp2*, with a significant decrease in PC1 and PC7, and an increase in PC9 and PC11 cells (Fig. 6A-B’, p < 0.05; two-sided unpaired t-test with Welch’s correction). While PC7 was similarly affected in *Foxp2*-cKO and *Foxp1/2*-cKO mice, PC1 exhibited a stepwise reduction, paralleling the gradual loss of the cerebellar hemisphere in these mutant mice, highlighting the essential role of PC1 in hemisphere development (Fig. 6B-C).

**Figure 6.**
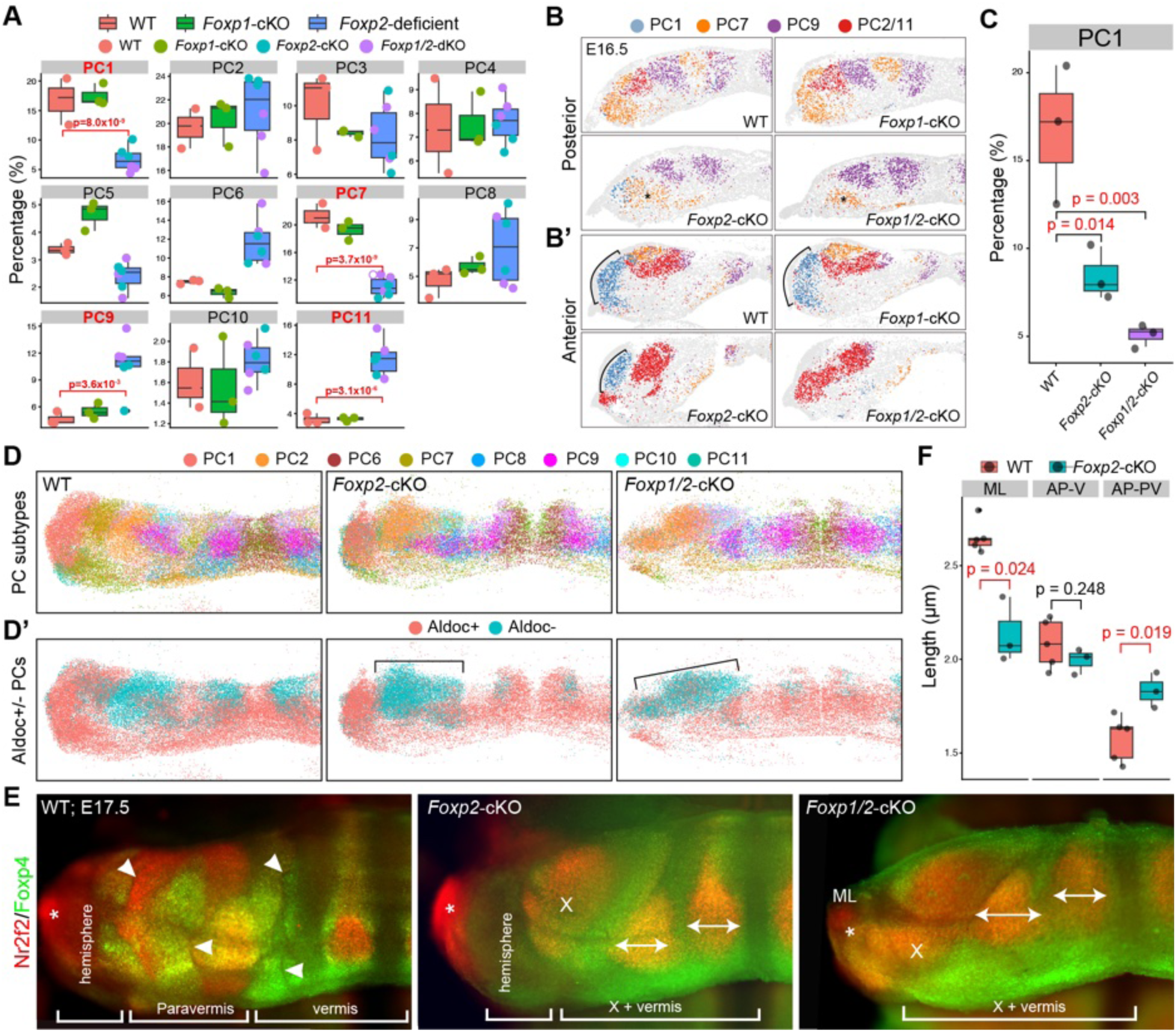
Regulation of PC diversification by *Foxp1* and *Foxp2*. (A) Quantification of PC subtypes across four genotypes (n = 3 per genotype) at E16.5. Significance p-values were calculated using two-way factorial ANOVA (*F*_(10, 110)_ = 18.66, *p* < 2e-16) and Tukey’s HSD post-hoc test. (**B** and **B’**) Significantly affected PC subtypes are shown in coronal sections at representative posterior (B) and anterior (B’) positions. Brackets denote PC1; asterisks indicate the reduction of PC7 in *Foxp2*-cKO and *Foxp1/2*-cKO cerebella. (**C**) Box plot showing stepwise reduction of PC1 in *Foxp2*-cKO and *Foxp1/2*-cKO cerebella. (**D** and **D’**) Representation of PC subtypes and predicted Aldoc^+/-^ identities in 3D reconstructed E16.5 cerebella. Brackets demarcate the single broad Aldoc^-^ bands flanking the vermis in *Foxp2*-deficient cerebella. (**E**) Volume images of E17.5 wholemount cerebella. Arrowheads indicate PC-sparse gaps; asterisks indicate Nr2f2^+^ cells outside the cerebellum; double arrows show two enlarged PC9 clusters that become much closer to each other in *the Foxp2*-deficient cerebella. (**F**) Measurements of circumference along the mediolateral axis (ML), anteroposterior axis in the vermis (AP-V), and anteroposterior axis in the paravermis (AP-PV). The p-values were calculated using the two-sided unpaired t-test with Welch’s correction.

To confirm the alterations in PC subtypes caused by the loss of *Foxp1* and *Foxp2*, we performed RNAscope FISH to examine additional markers that were associated with specific PC subtypes: *Zfhx4* for PC1, *Sulf1* for PC2, *Lmo4* for PC7, and *Sfrp1* for PC11. As expected, *Foxp1* transcripts were mostly absent from *the Foxp1*-cKO and *Foxp1/2*-cKO cerebella (Fig. S6A and B). *Zfhx4* marked PC1 cells and dentate nuclear neurons in *Foxp1*-cKO cerebella and revealed selective reduction and complete loss of PC1 in *Foxp2*-cKO and *Foxp1/2*-cKO cerebella, respectively (Fig. S6A). The expression of *Sulf1* and *Sfrp1* distinguished PC2 and PC11, respectively, and demonstrated an increase in PC11 cells in the *Foxp2*-deficient cerebella (Fig. S6A). *Lmo4* transcripts, which mark PC7, were absent in *Foxp2*-cKO and *Foxp1/2*-cKO mice (Fig. S6A). Notably, the combinatorial expression of *Lmo4*, *Ndst4*, and *Nrp2* revealed at least three PC1 subgroups, two of which were notably altered by the loss of *Foxp1*, suggesting a potential subdivision of this PC subtype (n = 4/4; Fig. S6B). Our data show that interactions between *Foxp1* and *Foxp2* are crucial for Purkinje cell diversification.

Using 3D reconstruction from serial sections and volume imaging, we corroborated the reduction in PC1 and PC7, as well as the expansion of PC9 and PC11 in the *Foxp2*-deficient cerebella (Fig. 6D-E). In the absence of *Foxp2*, the two PC9 clusters that resided in the vermis and paravermis were markedly expanded and became closer to each other, owing to the reduction in intervening PC clusters (Fig. 6B, D, and E). We applied scANKRS to transfer Aldoc^+^ and Aldoc adult PC identities onto the reconstructed 3D space in WT and mutant cerebella at E16.5 (Fig. 6D’). The resulting model correctly predicted the corresponding change in the longitudinal striped pattern in the adult cerebellum of *Foxp2*-cKO and *Foxp1/2*-cKO mice (Fig. 5F-H and 6D’). These findings further support the notion that topographic distribution of embryonic PC subtypes is predictive of alternating Aldoc^+/-^ PC stripes in the adult cerebellum.

At E17.5, several PC-sparse gaps disappeared in the prospective paravermis of the *Foxp2*-deficient cerebella (Fig. 6E), suggesting the loss of PC compartment boundaries. To examine whether this affects cerebellar cortex growth, we compared ML and AP lengths between the WT and *Foxp2*-cKO cerebella using 3D reconstructed images in Imaris for precise volumetric measurements. While the AP axis showed no significant difference in the vermis, it was significantly lengthened in the paravermis of *the Foxp2*-cKO cerebella (Fig. 6F). Conversely, the ML axis was significantly reduced in *the Foxp2*-cKO cerebella (Fig. 6F). These findings indicate that *Foxp1* and *Foxp2* cooperate to define PC subtype identities, and that proper PC diversification is essential for maintaining compartment boundaries and cerebellar cortex expansion.

### Importance of Foxp1-expressing PCs in the development and evolution of the cerebellar hemisphere

Given the critical role of PC1 in cerebellar hemisphere development, we examined PC1-like cells in the developing cerebella of humans and chickens, in which the cerebellar hemisphere is expanded and absent, respectively. Purkinje cells (PCs) were extracted from a published scRNA-seq dataset of the human fetal cerebellum ^32^ and aligned with the mouse PC subtypes. While similar PC heterogeneity was observed, the human cerebellum had significantly more abundant PC1 and PC4 but fewer PC6 (Fig. 7A). As PC1 and PC4 contribute to the lateral part of the cerebellum, and PC6 to the midline, these findings align with the enlargement of the hemisphere at the expense of the vermis in humans. As shown previously ^33^, *FOXP1* is uniformly expressed in PCs located in the lateral part of the fetal human cerebellum, which occupies a much larger region than the lateral cerebellum in mice (Fig. 7B and C).

**Figure 7.**
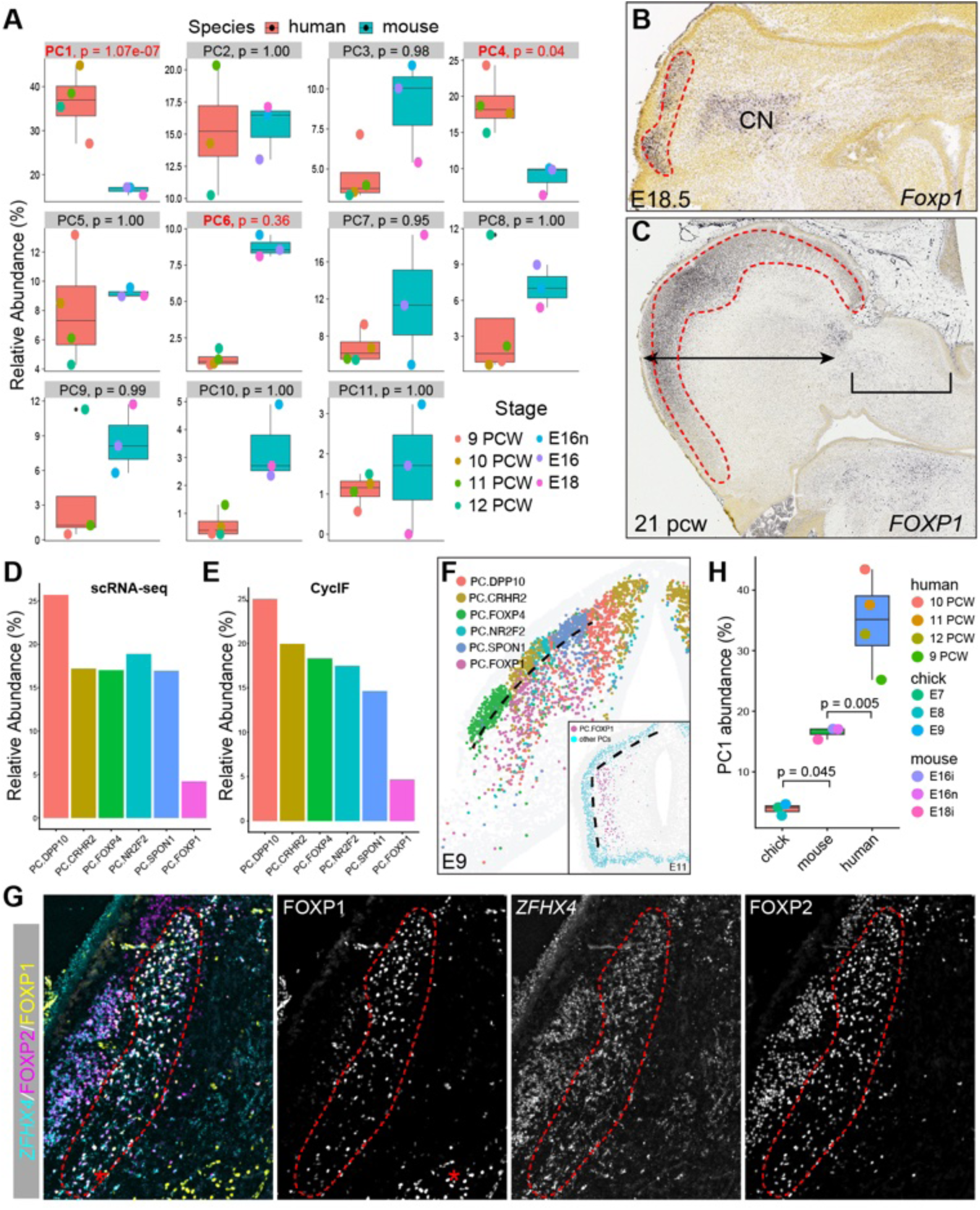
Divergent abundance of PC1 cells in the human, mouse, and chick cerebella. (**A**) Comparison of PC subtype abundance in the human and mouse cerebellum. (**B** and **C**) ISH on coronal sections of mouse (B) and human (C) cerebella. Dashed lines indicate *FOXP1*-expressing PCs in the cerebellar hemisphere. The bracket and double arrows indicate the vermis and the hemisphere, respectively. (**D** and **E**) Quantification of the abundance of different PC subtypes in E9 chick cerebellum using scRNA-seq and CycIF. (**F**) Representative images of the six PC subtypes in the coronal section of E9 chicken cerebellum. The dashed line indicates *FOXP1*-expressing PCs in the Purkinje cell layer (PCL). The inset shows that *the FOXP1*+ PC cluster remains beneath the PCL at E11. (**G)** IF for FOXP1 and FOXP2, and FISH for *ZFHX4* in the coronal section of the E9 chick cerebellum. Red dashed lines outline the FOXP1+/*ZFHX4*+ PCs beneath the PCL; asterisk indicates FOXP1 expressing CN neurons. (**F**) Box plots showing the abundance of PC1 across species. P-values were calculated using ANOVA (*F*_(2, 7)_ = 32.34, *p* = 0.0003) followed by Tukey’s post hoc test.

To investigate PC heterogeneity in the chick cerebellum, we conducted scRNA-seq of chick embryonic cerebella daily between E6 and E9 as PCs are born between E4 and E5.5 in the chick cerebellum ^34^. This produced a dataset of 13,064 cells, with a medium of 2,338 cells per stage (Fig. S7A and B). By combining scRNA-seq and spatial protein profiling, we identified six molecularly distinct PC subtypes organized as longitudinal stripes in the chick cerebellum at E9 and E11 (Fig. 7D-F and S7C). The least abundant PC subtype expressed *FOXP1* and *ZFHX4*, similar to PC1, in the mouse and human cerebellum (Fig. 7G and S6A). Therefore, there is stepwise increase of PC1 in the embryonic cerebellum from the chick, mouse, and to human, correlating the increase of the cerebellar hemisphere (Fig. 7H). Notably, PC1-like cells distinctively resided beneath the PC layer, which consisted of other five PC subtypes in the E9 and E11 chick cerebella (Fig. 7F and G). Pulse-chase labeling with 5-ethynyl-2’-deoxyuridine (EdU) revealed that PC1-like cells were born between E3.5 and E5.0, before the final wave of PC production at E5.5 (Fig. S7D-G). Therefore, the unique positioning of PC1-like cells is likely due to their inherent properties of cell migration and distribution rather than being a result of late birth.

Collectively, our data show that PC1 cells are significantly more abundant in the human cerebellum and less developed in the chick cerebellum. The progressive increase in PC1 abundance in the chick, mouse, and human cerebella suggests that this FOXP1^+^ PC subtype is pivotal for evolutionary expansion of the cerebellar hemisphere.

## DISCUSSION

In this study, we used scRNA-seq to systematically characterize PC heterogeneity and identified at least 11 molecularly distinct PC subtypes in the embryonic mouse cerebellum. By integrating scRNA-seq data from embryonic and adult stages with spatial expression profiling, we demonstrate that the 11 PC subtypes are confined to specific areas of the cerebellar cortex, predicting the regional specificity of PCs in adults. Additionally, we show that *Foxp1* and *Foxp2* are key regulators of PC subtype differentiation, with disruptions in these genes leading to altered cerebellar patterning. Notably, a *Foxp1*-expressing PC subtype plays a crucial role in cerebellar hemisphere development, offering insight into both cerebellar evolution and development.

The cerebellar cortex is organized into parallel circuit modules, each comprising distinct assemblies of PCs, inferior olivary neurons, and cerebellar nuclear neurons ^5,6^. A long-standing hypothesis suggests that parcellation of the cerebellar cortex is based on scaffolding provided by diverse PC subtypes ^5,6^. Fujita et al. identified 54 PC clusters in the prenatal mouse cerebellum, which were characterized by their position, shape, and expression of a few molecular markers ^35^. However, applying this classification in comparative analyses is challenging because of variability in sectioning planes between studies, especially when comparing cerebella with different morphologies between developmental stages or genetic backgrounds. By combining scRNA-seq and CycIF-based expression profiling, we established a workflow to map the topographic distribution of the PC subtypes identified by scRNA-seq. Importantly, this approach enables precise and efficient alignment of PC subtypes across different developmental stages, genetic conditions, and species. Notably, some PC subtypes (PC2, 3, 4, 7, 9, and 10) form two longitudinal bands on one side of the cerebellar anlage, displaying subtle differences in gene expression between these bands. For example, PC7 cells form longitudinal bands in both the paravermis and vermis, whereas Pcdh10 expression is only present in the lateral one in the paravermis (Fig. 3G). In the original classification proposed by Fujita et al., PC7 is subdivided into numerous clusters ^35^. Although this level of refinement is valuable, we believe that the current classification of the 11 PC subtypes offers sufficient granularity to facilitate robust comparative studies. Nevertheless, further subdivisions of certain PC subtypes are possible. Our data suggest that PC1 may contain subgroups, and future studies are needed to determine whether these subdivisions correspond to the lobulation of the hemispheres, which include the simplex, Crus I, Crus II, and pyramidal lobules – regions known to have distinct functions.

The development of longitudinal PC stripes is a gradual process involving dynamic changes in stripe markers, with most PC cluster markers downregulated before definitive stripes emerge ^36^. Linking embryonic PC clusters to adult PC stripes has been challenging, despite the known correlation between the timing of PC generation and their ML positioning ^37–39,39–41^. The longitudinal PC stripe pattern is refractory to experimental manipulations ^42–46^, and the mechanisms driving stripe development remain unclear. Here, we demonstrate that integrative scRNA-seq analyses can accurately predict adult PC phenotypes from embryonic subtypes (Fig. 3A-D). Perturbations in PC diversification due to *Foxp1* and *Foxp2* deletions altered PC stripe patterns as predicted (Fig. 6D and 6D’). Moreover, we show that PC subtypes exhibit regional specificity across cerebellar lobules (Fig. 3D). For instance, PC11 is associated with flocculus, and its increase corresponds to an enlarged flocculus in *the Foxp2*-deficient cerebella. These findings suggest that heterogeneous PC subtype boundaries align with future fissures along the anteroposterior axis.

Mutations in *FOXP1* and *FOXP2* are linked to overlapping developmental brain disorders, including autism spectrum disorders and speech impairments in humans ^47^. However, the mechanisms underlying these mutations and their primary target cell types in the brain remain unclear. Our findings show that removing *Foxp1* and *Foxp2* from the cerebellum results in vocalization defects and other behavioral changes in mice (Fig. 4C-E) ^28^, suggesting that disrupted cerebellar modularity may contribute to the neurological symptoms observed in humans with *FOXP1* and *FOXP2* mutations. *Foxp1* is selectively expressed in PC1, a subtype in the prospective cerebellar hemispheres, separated from adjacent subtypes (PC2 and PC7) by a PC-free gap. *Foxp2* deletion leads to a marked reduction in PC1 and a smaller hemisphere, while combined *Foxp1* and *Foxp2* deletion causes complete loss of both PC1 and the hemisphere. Notably, PC1-like cells are more abundant in the fetal human cerebellum than in mice and are rare in the chick cerebellum, underscoring the evolutionary importance of *FOXP1*-expressing PCs in cerebellar hemisphere development.

In summary, our study provides new insights into the spatiotemporal development of PC compartmentalization, which contributes to the formation of somatotopic areas in the cerebellar cortex. We propose that adult cerebellar compartments with similar molecular profiles, axonal projections, and functional localizations likely originate from a common early PC compartment. Further research is needed to clarify the molecular and cellular mechanisms driving cerebellar compartmentalization, and to explore the relationship between PC subtype heterogeneity and their specific connectivity patterns.

## MATERIALS AND METHODS

### Animals and tissue preparation

All animal procedures adhered to the guidelines of the University of Connecticut Health Center Animal Care Committee and the national and state policies. Mice were housed on a 12-hour light/dark cycle with ad libitum access to food and water. Genetically modified mouse strains, including *Atoh1-Cre* ^29^, *En1-Cre* ^48^, *Foxp1-flxoed* ^49^*, and Foxp2-floxed* ^23^, were obtained from Jackson Laboratory and maintained on a CD1 outbred background. Embryonic day (E) 0.5 was designated as the day a vaginal plug was detected. Primer sequences and PCR genotyping protocols are available on the JAX website. Fertilized chicken eggs were obtained from the University of Connecticut poultry farm and incubated at 38°C until the desired embryonic stages, with day one of incubation marked as E0. Both sexes were pooled for histological, scRNA-seq, and ultrasonic vocalization studies.

Mouse brains were dissected in ice-cold phosphate-buffered saline (PBS) and fixed in 4% paraformaldehyde (PFA) before P2. After P2, transcardial perfusion with PBS and PFA was performed before submersion fixation. The brains were cryoprotected, frozen in optimal cutting temperature (OCT) freezing medium (Sakura Finetek), and sectioned to 12-20 µm using a cryostat microtome (Leica CM3050S). The sections were mounted on ColorFrost Plus microscope slides (Fisher Scientific).

### Ultrasonic vocalization tests

For the USV test, the mouse pup was separated from its mother and littermates and placed in a sound-attenuating chamber with an ultrasonic frequency detector set to 60 kHz. Following a 60-second habituation period, the analysis commenced. Ultrasonic calls emitted over 6 minutes were recorded and analyzed for both number and duration using UltraVox XT 3.2 (Noldus).

### Single-molecule fluorescence in situ hybridization (smFISH)

Hybridization chain reaction was performed as previously described ^20^. Briefly, tissue sections were washed with PBS, equilibrated in hybridization buffer, and incubated overnight at 37°C with the probes. After hybridization, the slides were washed and equilibrated with amplification buffer. For HCR initiation, the hairpins h1 and h2 were snap-cooled, mixed, and applied to the slides. Following the HCR process, the slides were washed, mounted with SlowFade Gold Antifade mounting medium (Thermo Fisher Scientific), and imaged. All probes and hairpin oligonucleotides were purchased from Molecular Instruments.

### RNAScope HiPlex Assay

RNAScope HiPlex assay (Advanced Cell Diagnostics) was performed according to the manufacturer’s instructions. Briefly, slides were baked at 60°C for 30 min, fixed in 4% PFA for 15 min on ice, and dehydrated in a graded ethanol series (50%, 70%, and 100%). A 5-minute target retrieval step was conducted prior to protease treatment, which involved incubating the sections with protease III at 40°C for 15 min. Hybridization was carried out with 12 specific probes (catalog numbers: 485491-T1 (*Lmo4*), 866311-T2 (*Rreb1*), 495411-T3 (*Sulf1*), 500661-T4 (*Nrp2*), 473421-T5 (*Sorcs3*), 480601-T6 (*St6galnac5*), 313631-T7 (*Ndst4*), 531081-T8 (*Zfhx4*), 404981-T9 (*Sfrp1*), 586761-T10(*Foxp1*), 428791-T11 (*Foxp2*), and 442831-T12 (*Gpc5*) at 40°C for 2 hours, followed by three rounds of signal amplification. The nuclei were stained with DAPI. The detection of the 12 targets was accomplished in three rounds of imaging, with four targets visualized per round. Between each round, the fluorophores were cleaved and washed away to prepare for the next round of signal development.

### Cyclic immunofluorescence (CycIF)

Cryosections were thawed, air-dried overnight, post-fixed in 4% PFA, and permeabilized with Triton X-100. Antigen retrieval was performed using heated 0.01M sodium citrate buffer (pH6.0), followed by blocking with 10% normal donkey serum and incubation with primary and secondary antibodies. After washing, the slides were mounted in SlowFade™ Gold Antifade (Thermo Fisher) and imaged using a Zeiss Observer 7 microscope, visualizing 4-6 antibodies per round. For subsequent staining, antibodies were eluted with glycine-urea-guanidine hydrochloride buffer and the immunostaining cycle was repeated as needed. Antibodies used in this study are listed in Supplemental Table S3.

### Generation of multiplexed images

Raw images with multiple scenes were captured and stitched using Zeiss ZEN (v3.5) and then split into individual scenes in the czi format. Images from different staining rounds were registered using DAPI as a reference via Fiji *Linear Stack Alignment with the SIFT MultiChannel* plugin. DAPI images were projected onto a single 2D image and processed for nuclear segmentation using StarDist ^50^. The resulting nuclear masks were converted into ROIs and their spatial coordinates were recorded to map the cell positions. These ROIs were used to quantify the fluorescence intensities across the channels. The generated expression matrix, along with the spatial coordinates, was processed and analyzed in R.

### CycIF data analysis and integration with scRNA-seq

The CycIF-generated expression matrix was imported into Seurat ^51^ for cell clustering. Cells were filtered based on the nuclear size, shape, and intensity sum. Standard Seurat workflows were used for data normalization, clustering, and dimensional reduction. Seurat objects from different samples and genotypes were merged and jointly processed. Batch-effect correction was applied using Harmony ^52^ when needed. The Seurat functions *FindTransferAnchors* and *TransferData* were used to map the scRNA-seq clusters and transcriptome onto the CycIF data.

### Quantitative analysis of inferior olivary nuclei

Nuclear segmentation was performed using Fiji Stardist ^53^ based on DAPI staining. Manual corrections and marker identification for Foxp2 and Casp3 were performed using the Fiji *Annotater* plugin. The nomenclature and delineation of the inferior olivary subnuclei followed a previous report ^14^. Counts from the right and left sides of the inferior olive were averaged.

### Single-cell RNA-seq analysis

Single-cell suspensions were prepared as described previously ^17^, with sequencing libraries from E16.5 and E18.5 cerebella generated using Chromium v2 (10x Genomics). Additionally, a cerebellar library (E16.5) was created using Chromium v3. Reverse transcription and library generation were performed in accordance with the manufacturer’s instructions. Sequencing reads were demultiplexed and aligned to the mm10 mouse genome reference using CellRanger v3.1. Expression count matrices were generated using CellRanger count function. Count matrices were processed using the Seurat R package ^51^. Normalization, scaling, and identification of variable genes were performed using the Seurat *SCTransformation* function. The top 30 principal components were utilized for cell clustering, and uniform manifold approximation and projection (UMAP) was used to visualize the cell clusters. Seurat *FindAllMarkers* function was used to identify cluster-specific genes. Differential gene expression analysis was performed using the *wilcoxauc* function in the presto package ^55^. Cell type-specific genes were identified using the COSG package ^56^.

Sequencing libraries of the chick cerebellum were generated using the Evercode WT Mini Kit (Parse Bioscience). Sequencing reads were demultiplexed using split-pipe v1.0.6p and aligned to the chicken genome GRCg7b and Ensembl Release 110 annotations. The ‘split-pipe‘ pipeline was used to generate a cell-by-gene count matrix, which was processed with Seurat ^51^.

BAM files were obtained from the European Nucleotide Archive (PRJNA637987) and NCBI GEO (GSE118068) and then converted to FASTQ using bamtofastq v1.3.2. Expression count matrices were generated using the *cellranger count* function. Published scRNA-seq data were aligned with the in-house E16.5/E18.5 dataset using the Seurat *FindTransferAnchors* and *MapQuery* functions.

### Immunostaining and volume imaging of wholemount mouse cerebella

The detailed protocol for the procedure is available on protocols.io ^57^. Briefly, the cerebellum was dissected, meninges were removed, and the tissue was fixed in 4% PFA at 4°C for two hours, followed by three PBS washes. The samples were permeabilized with 0.05% Tween-20 and 0.2% Triton X-100 in PBS (TTP) at 37°C for 45 min and then treated with citric acid at 95°C in a steamer for 30 min. After blocking with 10% donkey serum in TTP, the samples were stained with primary and secondary antibodies for two days at 4°C. Clearing was performed using the modified RTF method ^58^. Samples were incubated at 4°C in the following solutions: 30% TEA, 40% formamide, and 30% H_2_O for 1 h; 60% TEA, 25% formamide, and 15% H_2_O for at least 4 h; and 70% TEA, 15% formamide, and 15% H_2_O for 2 h. For imaging and storage, the samples were immersed in 80% glycerol with the refractive index adjusted to 1.45, compatible with the Zeiss lightsheet Z1 microscope (5x objective). 3D reconstruction images and movies were generated using Imaris 9.0 (Bitplane).

### Volumetric analysis of the embryonic mouse cerebellum

To measure the volume of flocculus, intensity-based surface objects were generated by manually outlining the structure using the surface creation module in Imaris 9.0 (Bitplane). The measurement points were placed at the desired anatomical landmarks to assess the distance between the two points in the 3D space. The circumference of the structures was calculated by summing the distances between the sequential measurement points, effectively creating a polyline that represented the perimeter of the cerebellar cortical surface.

### Statistical analysis

Data processing, statistical analysis, and plotting were performed using R version 4.3.2. For comparisons between two groups, a two-sided unpaired t-test with Welch’s correction was applied, whereas ANOVA followed by Tukey’s HSD post-hoc test was used for comparisons involving more than two groups. No statistical methods were used to predetermine sample sizes. Sample sizes reflect those commonly used in the field and our previous studies. Normality and equal variance were not tested. No data or animals were excluded from this study. The results were reproducible for three or more samples, with quantitative data expressed as the mean ± SEM.

## ACKNOWLEDGMENTS

We thank Jihye Shin, Elliot Wilion, Kerry Morgan, Jasmine Aboumahboob, and Shravya Anisetti for their contributions to this study. We also thank Martina Ysabel Miranda for critically reading and providing valuable feedback on the manuscript. We are grateful to the JAX-UConn Single-Cell Genomics Center for their support with scRNA-seq experiments. We extend our thanks to Drs. Alex Joyner, Thomas Müller, and Bennett Novitch for providing the antibodies against En1/2, Tlx3, and Foxp1.

## AUTHOR CONTRIBUTIONS

Conception and supervision of the study: JL

Data generation: NF, QG, TP, RD, and JL

Data analysis: NF, QG, and JL

Data interpretation: NF, QG, and JL

Writing the manuscript: JL

Review and editing of the manuscript: NF, QG, and JL

## Data and material availability

Sequencing data were deposited in GEO under accession codes GSE256438 and GSE278923. The publicly available datasets were used in this study: GSE118068 and PRJNA637987.

## Code availability

Computer codes associated with the manuscript are available on the JLiLab GitHub: https://github.com/JLiLab/Studying-Parkinje-cell-heterogeneity

## SUPPLEMENTAL MATERIALS

**Supplementary Movie 1. 3D Rendering of PC-subtype cluster and stripe development.** 3D-rendered volume imaging of whole-mount mouse cerebella from E14.5 to P6, following immunofluorescence staining and tissue clearing. Dashed lines indicate PC-sparse gaps.

**Supplementary Table S1. Molecular features of major cerebellar cell types.** An excel table showing molecular features of cerebellar cell types.

**Supplementary Table S2. Molecular features and functional enrichment of different PC subtypes**. Excel tables showing molecular features (sheet 1) and GO-term enrichment (sheet 2).

**Supplementary Table S3. List of antibodies used in the study**. An excel table of antibodies used in this study.

---

**Figure S1.**
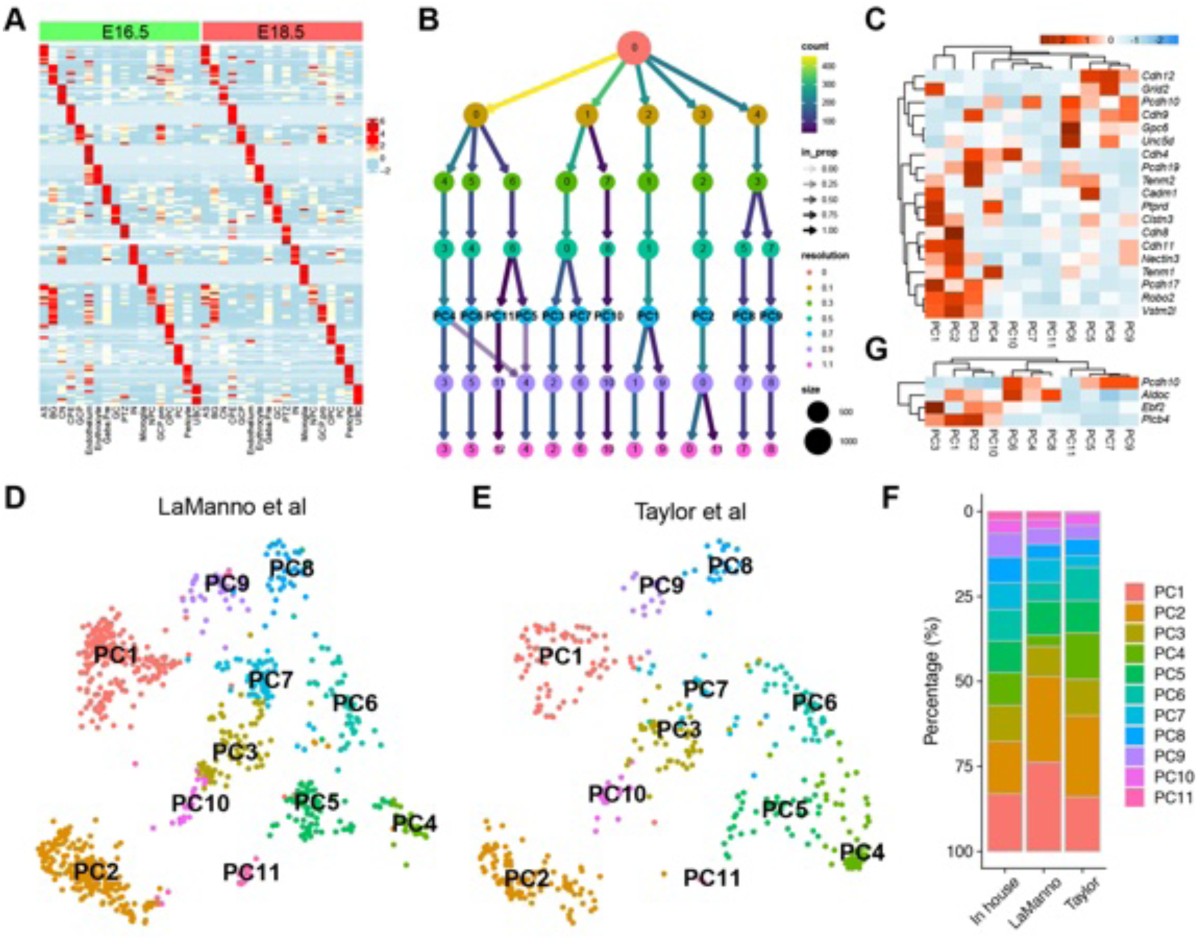
Evaluation of the robustness of clustering PC subtypes. (**A**) Heatmap showing similar molecular features of different cerebellar cell types at E16.5 and E18.5. (**B**) Cell clustering results with an increasing resolution from 0 to 1.1. PCs are gradually resolved from one to 11 groups as the resolution increases from 0 to 0.7, after which only minor changes are observed. (**C**) Heatmap showing the differential expression of cell adhesion molecules in the plasma membrane among PC subtypes. (**D** and **E**) Identification of similar 11 PC clusters in two published scRNA-seq datasets of E16.5 and E18.5. (**F**) Bar plots showing the relative abundances of different PC subtypes in the three independent scRNA-seq datasets. (**G**) Heatmap showing the differential expression of known Aldoc+/-markers among PC subtypes.

**Figure S2.**
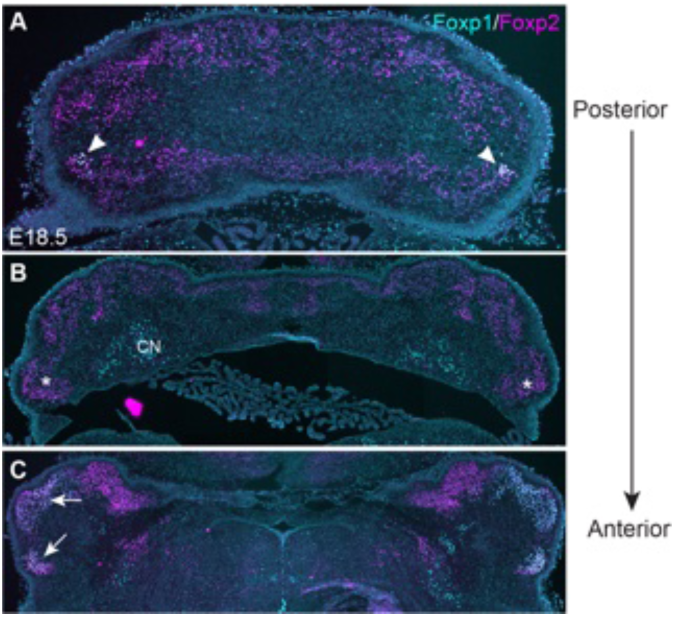
*Foxp1*-expressing PCs occupy the lateral-most portion of the cerebellum. (**A-C**) Immunofluorescence (IF) for Foxp1 and Foxp2 in three coronal sections of the E18.5 cerebellum corresponding to different AP positions. Arrowheads and arrows indicate *Foxp1*^+^ PCs in the lateral end of lobe X and the hemisphere, respectively. The asterisk denotes the absence of Foxp1^+^ PCs in the intervening section.

**Figure S3.**
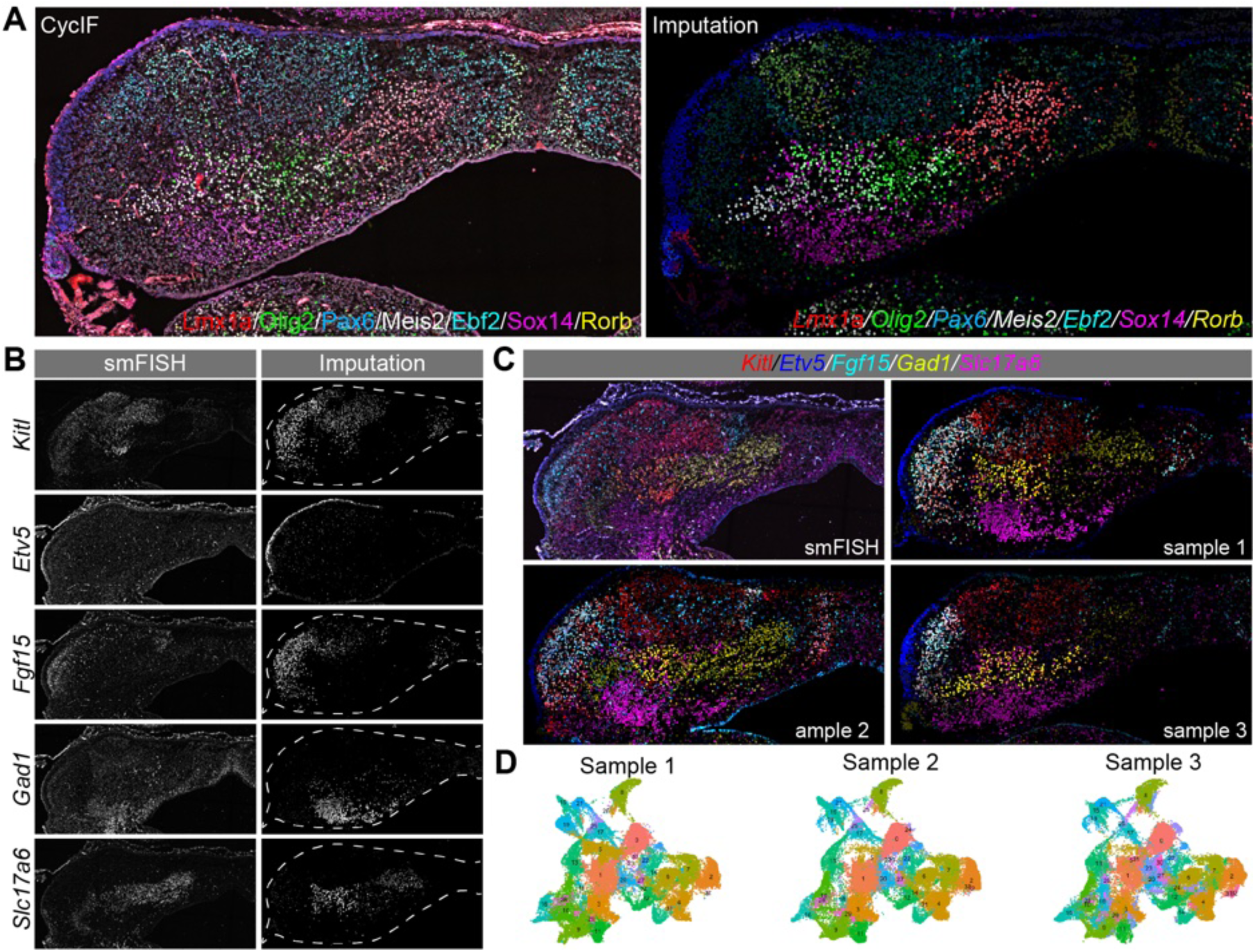
Evaluation of accuracy and reproducibility of scANKRS. **(A)** Comparison of cyclic immunofluorescence and predicted gene expression. **(B** and **C)** Expression of selected genes based on single-molecular fluorescence in situ hybridization (smFISH) and scANKRS-based imputation in single channels (C) and merged colors (D) in a coronal section of three E16.5 cerebella. **(D)** UMAP representation of cell clusters according to the transcriptome imputed from three independent experiments.

**Figure S4.**
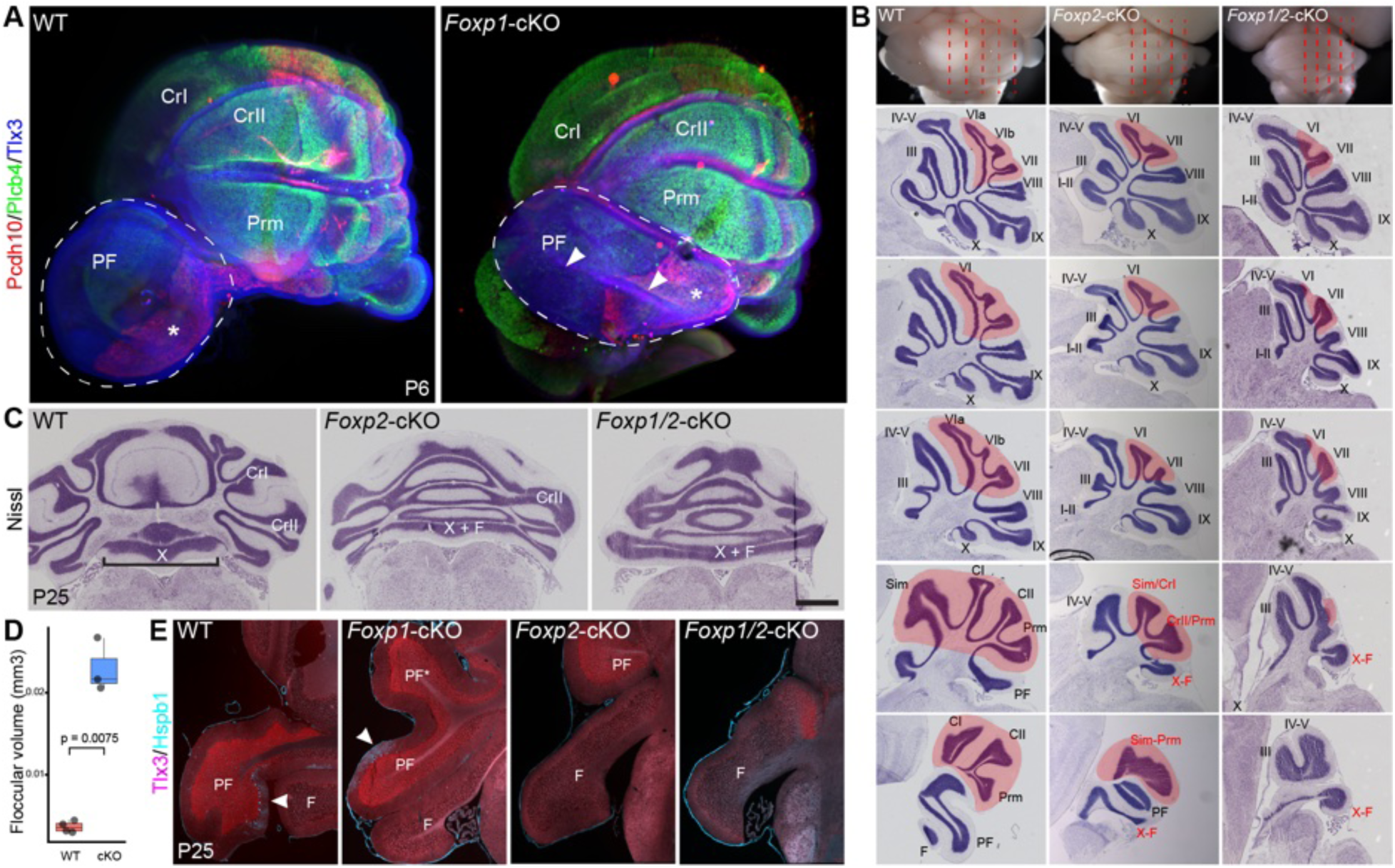
Abnormal cerebellar development without *Foxp1* and *Foxp2*. (**A**) LSFM images of P6 cerebella. Paraflocculus is outlined by dashed lines. The asterisk indicates the Pcdh10+ domain in the posterior part of the paraflocculus, and the arrowheads denote the abnormal fissure. (**B** and **C**) Nissl histology of the sagittal (B) and coronal (C) cerebellar sections at P25. The central lobe is highlighted in pink. The names of severely affected lobules are shown in red. Note that lobule X is restricted to the vermis (bracket) in the WT cerebellum, but becomes continuous with the flocculus without *Foxp2*. (**D**) Volumetric measurements of the flocculus in E17.5 cerebella. The p-value was calculated using the two-sided unpaired t-test with Welch’s correction. (**E**) Immunofluorescence for Tlx3 and Hspb1 in coronal sections of the P25 cerebella. Arrowheads indicate Hspb1 immunoreactivity in the paraflocculus.

**Figure S5.**
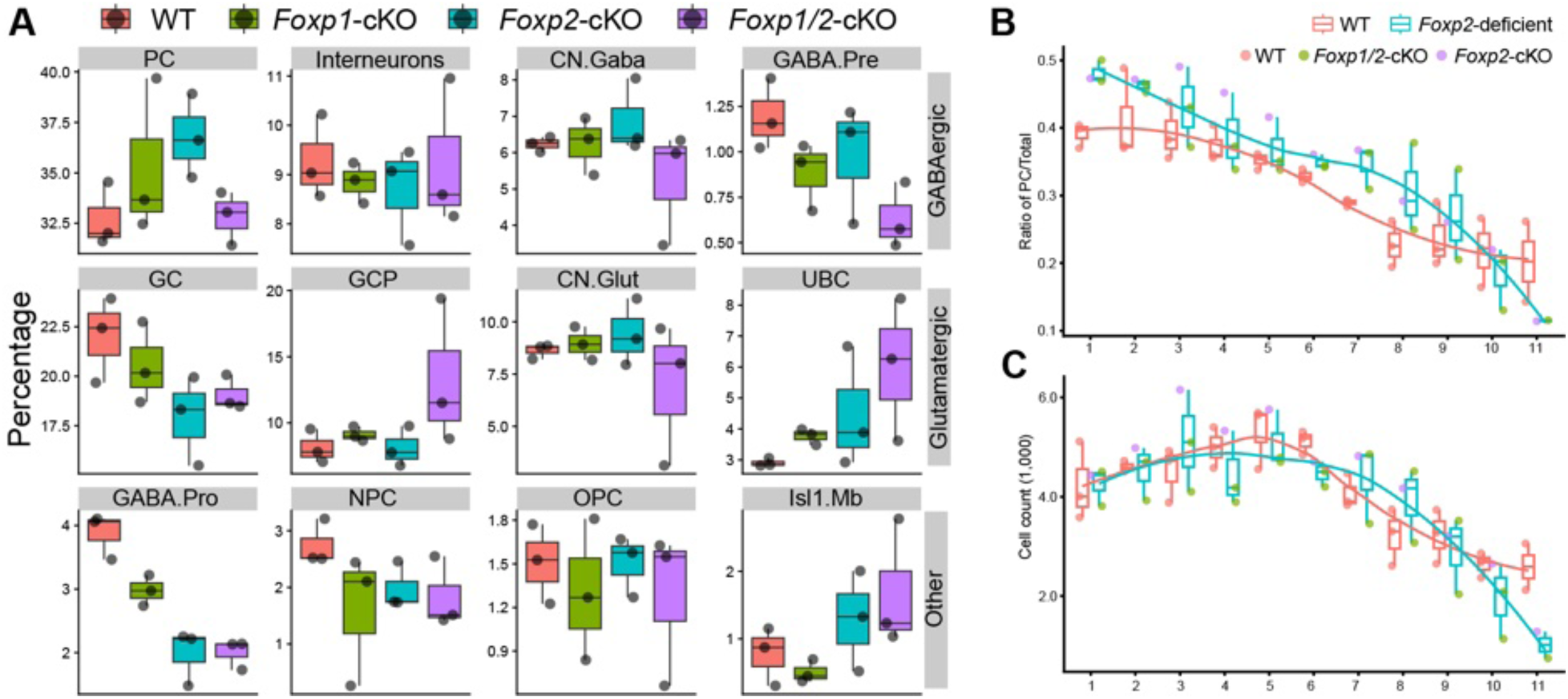
Loss of *Foxp1* and *Foxp2* has little impact on the relative abundance of major cerebellar cell types, including PCs. (A-C) Box plots showing the relative abundance of major cerebellar cell types (A and B) and the total number of PCs. The numbers on the X-axis represent serial coronal sections from posterior to anterior. No statistically significant differences were observed between the WT and mutants. ANOVA for A: significant effect of cell type [F_(7, 64)_ = 496.22, *p* < 2e-16] but no main effect of genotype [*F*_(3, 64)_ = 0.00, *p* = 1.000]. The interaction between cell type and genotype was not significant [*F*_(21, 64)_ = 1.60, *p* = 0.079)); two-sided unpaired t-test with Welch’s correction for B.

**Figure S6.**
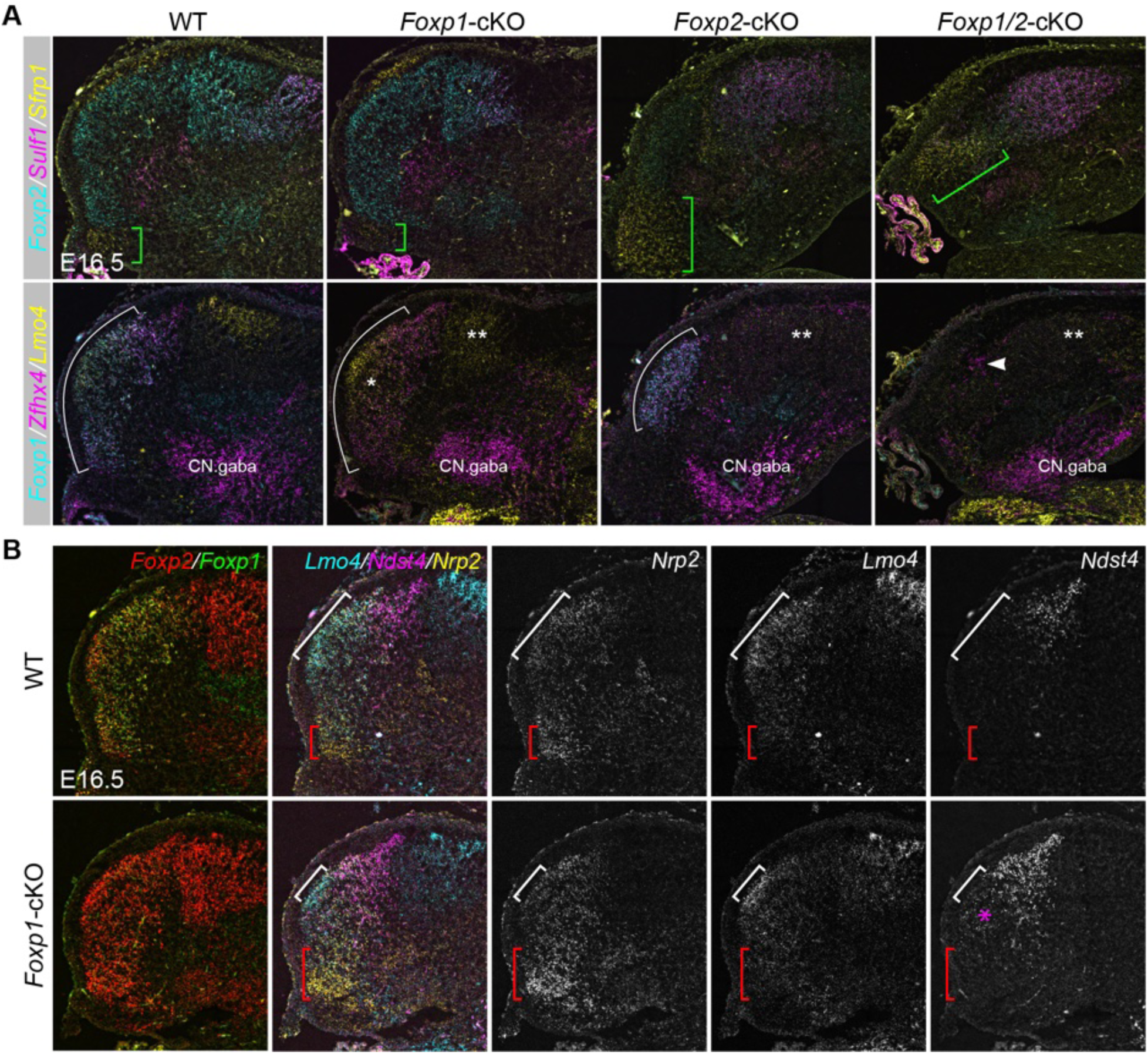
Validation of altered PC subtype abundance due to the loss of *Foxp1* and *Foxp2*. (A, B) RNAscope HiPlex assay of coronal sections from E16.5 cerebella of the indicated genotypes. In (A), white and green brackets indicate PC1 and PC11, respectively. White and green brackets in (A) indicate the boundaries of PC1 and PC11, respectively. A single asterisk (*) indicates ectopic expression of Lmo4, while double asterisks (**) denote the loss of Lmo4+ PC7 in the *Foxp2*-cKO and *Foxp1/2*-cKO cerebella. In (B) white and red brackets highlight two PC1 subdomains that are enlarged or shrunken in the *Foxp1*-cKO cerebellum.

**Figure S7.**
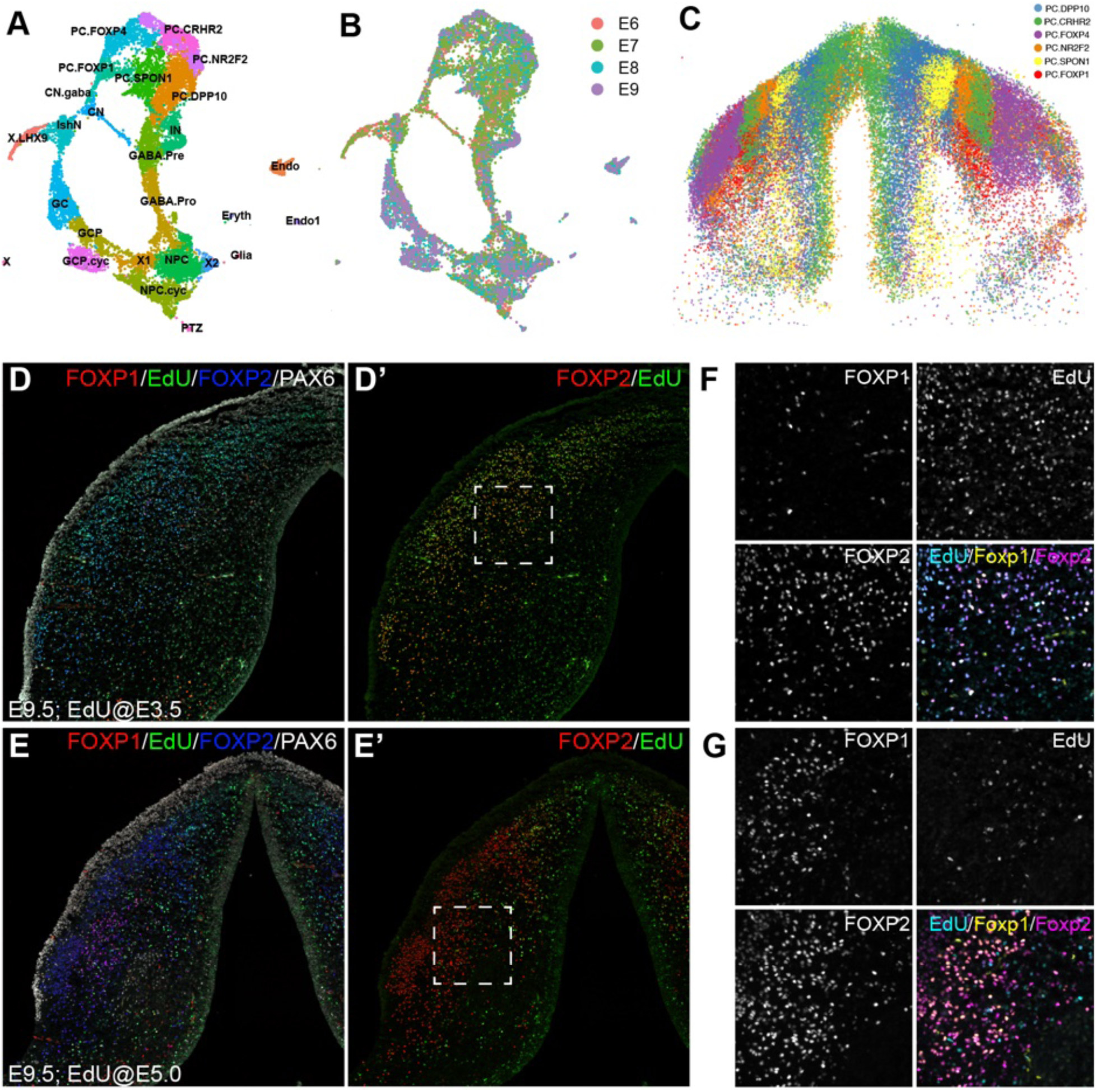
scRNA-seq and birth-dating analysis of the chick cerebellum. (A and B) UMAP projection of scRNA-seq data from embryonic chick cerebellum between E6 and E9. (C) Three-dimensional reconstruction of six Purkinje cell (PC) subtypes in the chick cerebellum at E9 based on CycIF analysis. (D-G) Immunofluorescence and EdU pulse-chase assays in coronal sections of the E9 chick cerebellum. The boxed areas in D’ and E’ are shown at higher magnifications in F and G.

